# Mapping prion pathology in mice using quantitative imaging: an MRI study

**DOI:** 10.1101/2021.12.28.474373

**Authors:** Eleni Demetriou, Mohamed Tachrount, Matthew Ellis, Jackie Linehan, Sebastian Brandner, John Collinge, Simon Mead, Karin Shmueli, Mark Farrow, Xavier Golay

**Affiliations:** Brain Repair & Rehabilitation, Institute of Neurology, University College of London, United Kingdom; Wellcome Centre for Integrative Neuroimaging, FMRIB, Nuffield Department of Clinical Neurosciences, University of Oxford; Department of Neurodegenerative Diseases, UCL Queen Square Institute of Neurology, London, United Kingdom; MRC prion unit, Institute of Neurology, London, United Kingdom; Medical Physics and Biomedical Engineering, University College of London, London, United Kingdom; MRC Prion unit, Institute of Neurology, London, United Kingdom

**Keywords:** MTR, MRS, T_1_, T_2_, Prion disease, MRI

## Abstract

Human prion diseases are fatal neurodegenerative disorders which cause cognitive impairment and neurological deficits. Additional measures of tissue status are necessary for improving the sensitivity and specificity of clinical diagnosis as in many cases clinical forms of prion disease are commonly mistaken for other forms of dementia. To that effect, we developed a set of quantitative magnetic resonance-based tools, including magnetic resonance spectroscopy (MRS), magnetization transfer ratio (MTR) and quantitative T_1_ and T_2_ imaging to study the course of the disease in an animal model of prion disease. Using *in vivo* MTR, significant changes were detected in the cortex and thalamus of late-stage prion -infected mice as compared to littermates. In addition, we found a significant increase of MTR in thalamus and cortex of 80 dpi healthy mice when compared with 160 dpi healthy mice suggestive of changes occurring during the development of the brain. Using quantitative T_2_ mapping, significantly higher values were measured in thalamus of prion mice at all stages of the disease (T_2_=40ms) while T_1_ was found to be significantly higher in cortex (T_1_=1.89s) and hippocampus, albeit only in late-stage prion mice as compared to aged-matched controls (T_1_=1.67s). Using quantitative MRS significant changes were detected in glutamate (Glu) and myo-inositol (Ins) at all stages of prion disease when compared with the control group. NAA, Cr, Lactate and Lipids were only found to be significantly different at early and late stages of the disease while Taurine (Tau) was only significantly increased in the asymptomatic stage without any significant change at early and late stages of the disease. These changes in MRI and MRS signals, which precede clinical signs of disease, could provide insights into the pathogenesis of this disease and may enable early detection of pathology.

## Introduction

Prion diseases are fatal neurodegenerative disorders with clinically silent incubation periods until the onset of the disease. Human forms of prion disease include sporadic and familial Creutzfeldt-Jakob disease (CJD) affecting one to two people per million population worldwide annually, whilst for genetic forms such as Gerstmann-Straussler-Scheinker (GSS) (1) syndrome and Fatal Familial Insomnia (FFI) (1), the incidence is ten times less. The neuropathogenesis of prion disease is strongly associated with the conversion of normal synaptic prion protein to the protease resistant forms termed PrP^Sc^. These are fatal diseases with no treatment reported so far. Early pathological aspects have been defined experimentally, but are difficult to translate into clinical practice. Imaging methods are widely available and refined imaging sequences for early diagnosis of CJD would be highly desirable to complement existing biochemical tests such as determination of 14-3-3 protein in the CSF (2).

To date, numerous models of prion disease have been developed in mice mimicking various histological features such as accumulation of misfolded prion protein, vacuolation, neuronal cell death, astrocytosis and microglia activation (3). In these models the length of the incubation period varies while developed clinical signs include ataxia and abnormal posture. The state of the art of imaging in prion disease targets deep and cortical grey matter structures using either hyper intensities in *T*_2_-weighted (*T*_2_w) imaging or microstructural changes in DWI scans (4). In hamsters, increased signal intensity on *T*_2_-weighted images was observed in the hippocampus at a pre-symptomatic stage (120 days) with the signal alterations being more pronounced with disease severity. The observed neuropathological changes included accumulation of PrP□□ and astrogliosis without any spongiform changes (5). In a second study, in which sheep were exposed to Prion disease, MRI findings included diffuse cerebral atrophy in asymptomatic and symptomatic sheep, with the origin of this finding being unclear (6). Although preclinical studies remain limited, the findings in animal models agree with clinical studies of prion disease, without, however, direct translatable biomarker to date being developed. In addition, the majority of imaging studies focused on identification of structural biomarkers for monitoring disease progression. Molecular MRI provides an additional level of information, thus, extending imaging parameters beyond the anatomical level. Moreover, it holds the potential for identification of early stages of prion disease since metabolic changes have been shown to precede any structural alterations (7).

The primary aim of this study was to identify early translational biomarkers of prion disease that can be detected non-invasively by MR. In the current study, we set out to develop a range of quantitative MRI techniques for the mapping of T_1_ and T_2_ as well as Magnetization transfer (MT) in addition to MRS to investigate biochemical changes that occur longitudinally in (Rocky Mountain Laboratory) RML mice intracerebrally inoculated with misfolded prion proteins. We hypothesised that MRS could provide a general read out of metabolic changes in the brain of prion mice which are then combined with quantitative structural biomarkers. Any observed changes will be correlated with well described histopathological features that underpin disease progression.

## Methods

### Animal studies

#### Control and prion-infected mice

Work with mice was performed under licence granted by the UK Home Office and conformed to institutional guidelines. Two groups of 7-week-old RML mice were intracerebrally inoculated with 30μl of 1% brain homogenate from RML prion-infected mice (n=19) or brain homogenate from uninfected mice as controls (n=11). The prion-infected group was separated into three groups of mice scanned at different stages of prion disease: 80 days post injection (dpi) – asymptomatic-stage (n=6), 130 dpi – early-stage (n=6), and 160 dpi – late-stage (n=7). Control mice were separated into two groups: 80 dpi (n=5) and 160 dpi (n=6). All mice were anaesthetized (1.5-1.8% isoflurane in 1.5 l/min air with balance in oxygen) and scanned on a 9.4 T Agilent system (Agilent Technologies, Santa Clara, CA, USA) using a 33-mm-diameter transmit/receive coil (Rapid Biomedical GmbH, Rimpar, Germany).

#### Anatomical MRI and MRS protocol

T_2w_ anatomical scans were acquired in an axial slice using a fast spin-echo sequence (data matrix: 256×128, TR=3000ms, TE=20ms, FOV=20×20mm). *In vivo* proton spectra were acquired using a point resolved spectroscopy (PRESS) sequence (TR = 5000msec, TE = 7.5msec, 128 averages, total acquisition time=10 min) with a voxel centred on the thalamus (1.7 × 4.3 × 1.8 mm□). After first and second order shimming, the typical linewidth of the water resonance was 20-23Hz.

##### MRS data analysis

The acquired spectra were processed and quantified using TARQUIN (8). Experimental data were modelled as a linear combination of modified simulated basis signals. The Cramer-Rao lower bounds were used as a reliability measure of the metabolite concentration estimates (below 20%). All metabolite concentrations are presented as mean ± standard deviation. Statistical analysis was performed using a two-tailed t-test. Significant changes in metabolite concentrations are indicated by p<0.05.

#### Magnetization transfer (MT) protocol

All images were acquired in a single slice (thickness=2mm), centred on thalamus. MT measurements were acquired using a gradient-echo sequence (matrix: 64×64, TR=2.11ms, TE=1.07ms, FOV=20×20mm^2^) with a train of off-resonance Gaussian pulses applied at 10 ppm and with an irradiation amplitude of 10μT (n=30, pulse length=50ms, flip angle (FA) =6000°, 99% duty cycle). We used a relatively high amplitude of the irradiation pulse for maximising the MT weighting. In addition, for improving the image SNR and reducing the physiological noise during MRI acquisition the measurements were collected three times and the averaged values have been calculated for each mouse.

#### MT Data analysis

In this study, MTR was calculated as (M_sat_ (10 ppm) - M_0_)/M_0_ where M_0_ is the signal without the application of irradiation RF pulses and M_sat_ (10 ppm) corresponds to the signal measured at 10ppm after the application of magnetization transfer pulses. Regions-of-interest (ROIs) were drawn on the anatomical image of each mouse in the cortex and thalamus for analysis of the corresponding MTR maps.

#### T_1_ and T_2_ measurements

An inversion recovery echo planar imaging (EPI) sequence was used to quantify the T_1_ values. A global adiabatic inversion pulse (FA=180°, duration=2ms) and 10 inversion times exponentially spaced were applied from 8.1ms to 7.5s. For the quantification of T_2_ values, a (Carr-Purcell-Meiboom-Gill) CPMG sequence consisting of a 90° Sinc-shaped RF excitation pulse along the x axis (duration=2ms) was used, followed by 15 Sinc-shaped refocusing RF pulses along the y axis (fa=180°, duration=1.6ms).

#### Post processing

T_1_ and T_2_ maps were obtained using custom-written scripts in MATLAB (Mathworks Waltham, MA) and by assuming mono-exponential decay for longitudinal and transverse relaxation.

#### Statistical analysis

Differences in metabolite’s concentrations among control animals, asymptomatic animals, mice showing early signs of disease and late-stage prion infected mice were assessed using mixed-effects regression models in STATA (9). In contrary to a simple t-test, a mixed-effects analysis allows the examination of multiple variables simultaneously for predicting an outcome measure of interest. Significant changes detected based on a fixed effect multivariate analysis are more robust because the whole group of prion mice is compared to controls based on a group of parameters rather than individual comparisons. The relationship between metabolite concentrations, T_1_, T_2_, MT and histopathological scores was assessed across the whole group of prion infected mice with Spearman’s rank correlation.

A detailed description of the quantification of histological biomarkers in prion-infected mice such as misfolded prion protein accumulation, astrogliosis, microglia activity and spongiosis is detailed in our previous studies (10). Here we will only present significant correlations with between imaging results and histology for the two groups of diseased and healthy mice. Finally, MRS data were correlated with our previously published (10) CEST results for assessing any correlations between metabolites and CEST signal. A value of p <0.05 was considered significant.

## Results

To investigate imaging abnormalities in prion-infected mice experimental work was carried out to assess whether significant changes in metabolites concentration, or alterations in MTR could be detected between prion-infected and control mice. Additional measures of tissue status were evaluated by means of quantitative T_1_ and T_2_ maps.

### MRS studies

The neurochemical profile of prion disease in mice at different disease stages was evaluated using high-quality MR spectra obtained in thalamus. Figure 1 shows representative MRS spectra in thalamus from a control and a prion-infected mouse at 160 dpi. The seven most abundant metabolites in the brain with CRBs < 20% were measured *in vivo* longitudinally. The metabolite concentrations were evaluated for age dependence between the two control groups scanned at 80 dpi and 160 dpi. The only change with age between both control groups was an increase in N-acetylaspartate (NAA) (p=0.009) and creatine (Cr) (p=0.005). Therefore, the control groups were merged into one group when compared with prion-infected mice for all the metabolites except NAA and Cr. Significant changes were detected in glutamate (Glu) and myo-inositol (Ins) at all stages of prion disease (80 dpi, 130 dpi, 160 dpi) when compared with the control group. However, there was no significant change in Choline (Cho). NAA, Cr, Lactate and Lipids were only found to be significantly different at 130 dpi and 160 dpi compared with the control group. Moreover, Taurine (Tau) was only significantly increased at 80 dpi without any significant change at 130 dpi and 160 dpi (see Figure 2). A mixed effects model analysis further confirmed the significant changes found with independent t-tests. Glutamate (Glu), Myoinositol (mI), lactate and lipids were found significantly different (p<0.05) at all stages of the disease when compared with healthy mice. Glutamate was correlated with astrogliosis (R^2^=0.81) and both lactate and lipids were also correlated with astrogliosis with R^2^=0.71. NAA was correlated with both astrogliosis and misfolded prion protein deposition with R^2^=0.6. No other significant correlations were found for the remaining metabolites.

**Figure 1.**
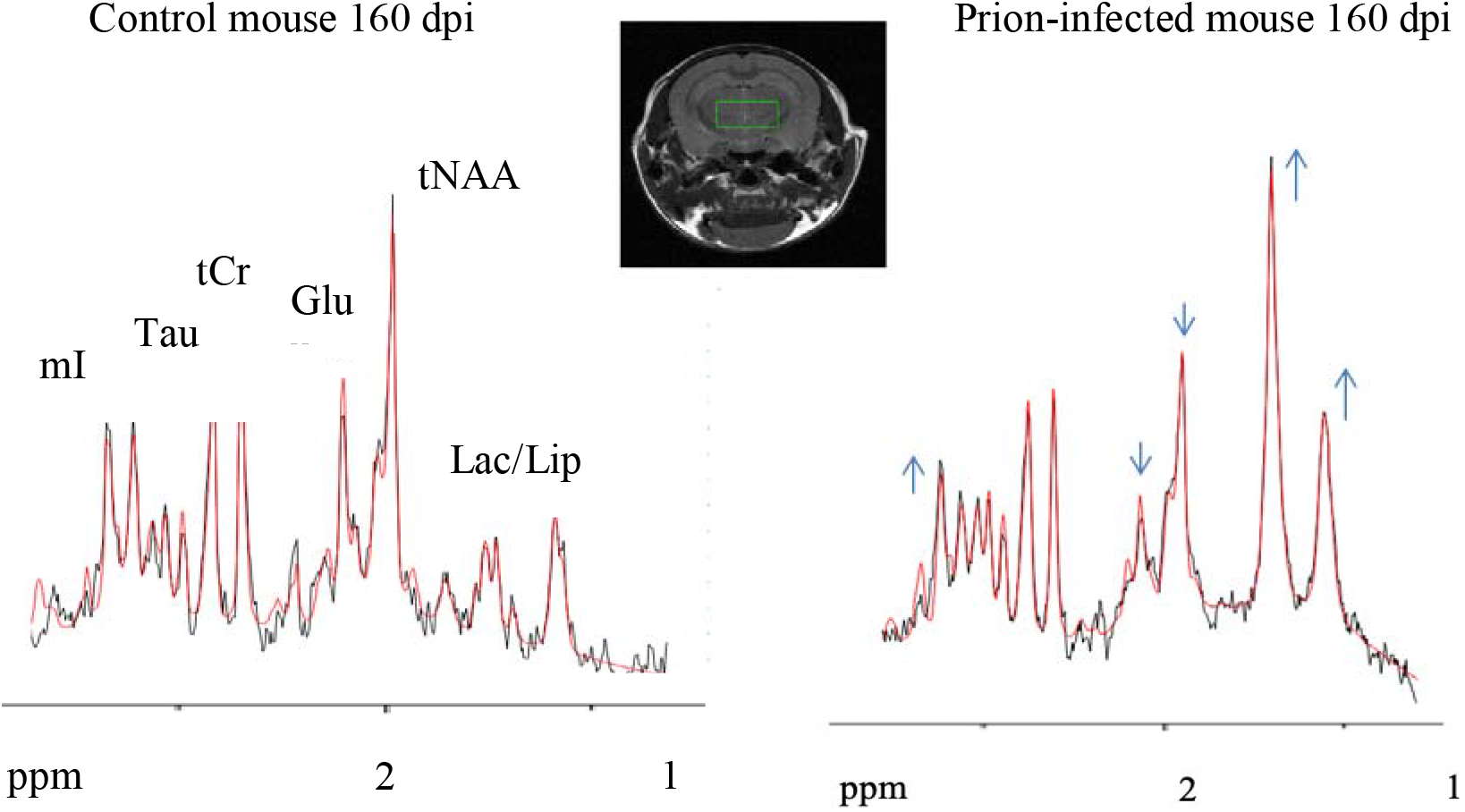
Representative ^1^H MRS spectra acquired in thalamus of a prion-infected mouse (b) and a control mouse (a) at 160 dpi. The arrows indicate metabolites that showed significant differences in concentration between the prion-infected and control groups. The PRESS voxel is in the thalamus is shown overlaid on a representative anatomical image.

**Figure 2.**
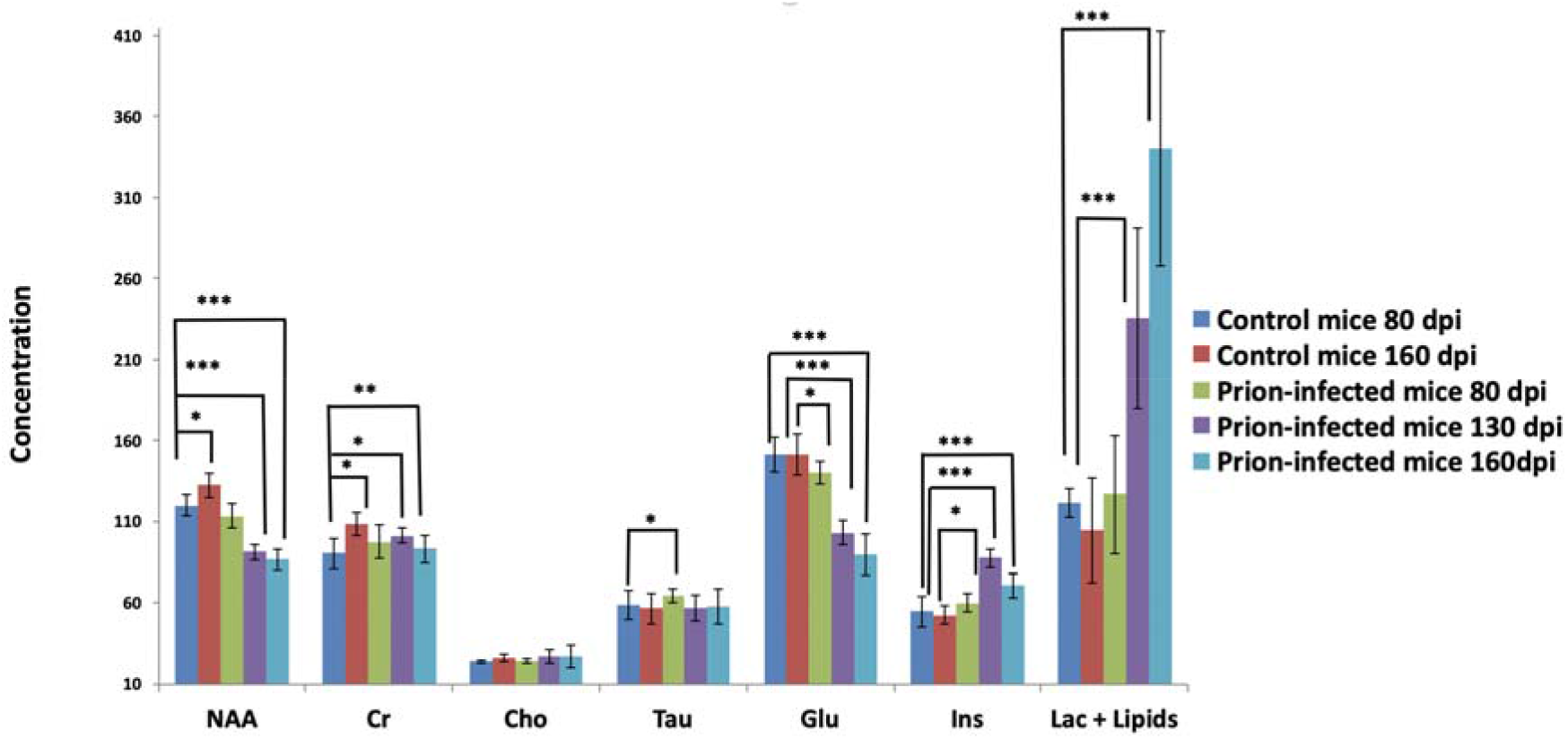
Comparison of the concentration of metabolites in thalamus between prion-infected mice at different stages of Prion disease and control mice. Data are expressed as mean ± standard deviation (* p<0.05, ** p<0.005, *** p<0.0005).

**Figure 3.**
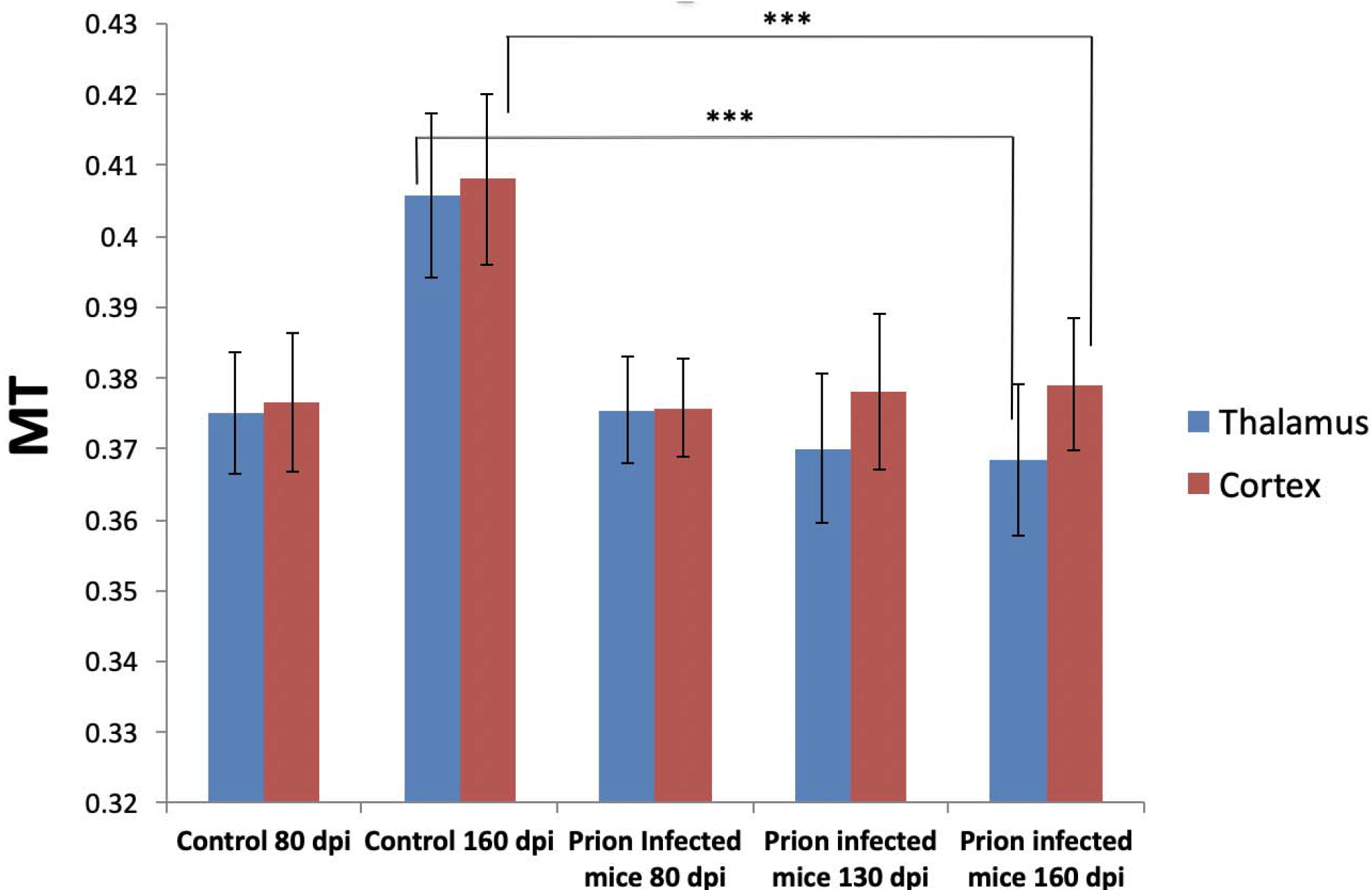
MTR at 20kHz in thalamus and cortex for prion-infected mice and control mice. Two-tailed t-tests were used to evaluate changes in MTR values measured in prion-infected and control mice (*** p < 0.001). A significant reduction in MTR ratio has been observed in thalamus and cortex of 160 dpi prion infected mice when compared with 160 dpi control mice. A significant increase in MTR has been observed for the 160 dpi control mice when they were compared with 80 dpi control mice.

### Correlations of CEST metrics with MRS

The MTR asymmetry at 3.5 ppm (10) was correlated with NAA (R^2^=0.58), with glutamate (R^2^=0.43), and with lactate and lipids (R^2^=0.64). The CEST signal ranging between 1-2 ppm was correlated with Myoinositol (R^2^=0.41). Finally, the MTR asymmetry ranging between 2-3 ppm was correlated with lactate and lipids (R^2^=0.51).

### Magnetization transfer results

MTR was evaluated to visualise changes in macromolecular concentration and composition, which might take place through both the developmental changes of the brain due to age difference between the mice groups as well as through the direct effect of the disease. Significant changes were detected in the cortex and the thalamus of 160 dpi prion-infected mice when compared to 160 dpi control mice (see Figure 4). Moreover, a significant increase of MTR was detected in thalamus and cortex of 160 dpi versus 80 dpi control mice. The MTR levels for the early and late-stage prion-infected mice were not significantly different, however, control mice of 130 days old were not scanned therefore it is difficult to conclude if there are any significant MT changes at the early stage of the disease. No significant correlations were detected between MT and histopathological markers or between MT and CEST results (10).

**Figure 4.**
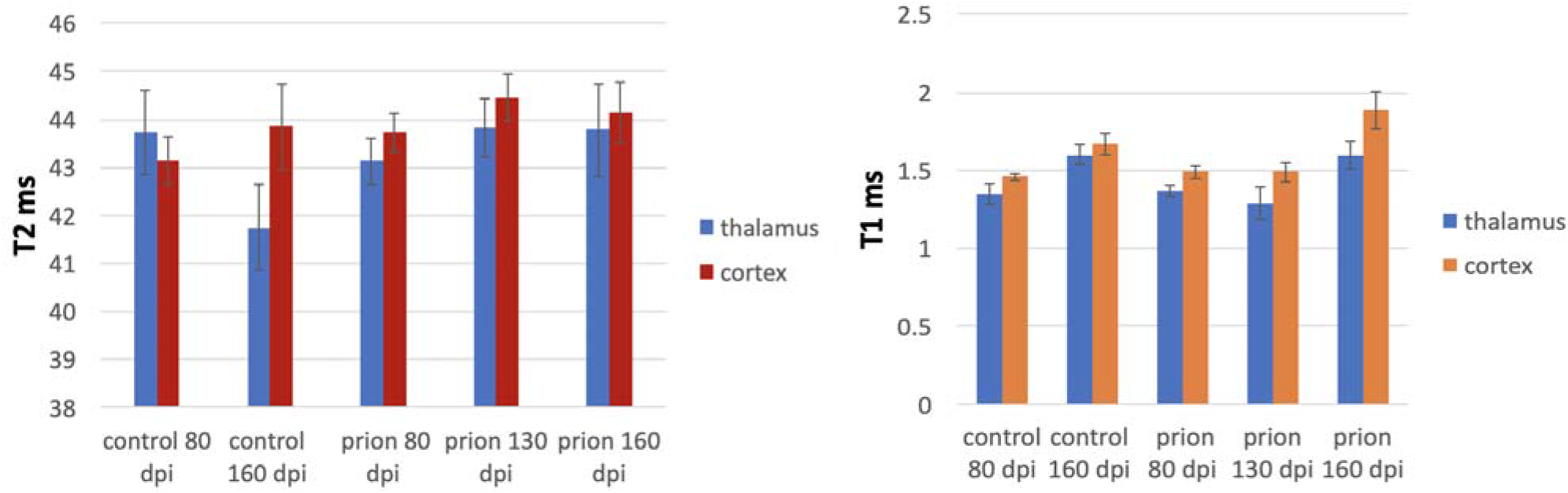
T1 and T2 in thalamus and cortex for prion-infected mice and control mice. A significant reduction in T_2_ mapping, were measured in thalamus of prion mice at all stages of the disease while T_1_ was found to be significantly higher in cortex, only in late-stage prion mice as compared to aged-matched controls.

### T_1_ and T_2_ results

T_2_ values were found to be significantly higher in thalamus of prion mice at all stages of the disease (T_2_=40ms, p=0.001) when compared to control mice (T_2_=37ms) while the T_1_ was found to be significantly higher in cortex (T_1_=1.89s, p= 0.0007) and hippocampus of only late stage prion-infected mice when compared to age-matched controls (T_1_=1.67s) (see figure 4). No significant correlations were detected between T_1_/T_2_ and histopathological markers or between our previously published CEST results and T_1_/T_2_ (10).

## Discussion

In this study, we used quantitative MRI measures for understanding the preclinical development of prion disease and assess which one would be easiest to translate into clinical practice. Indeed, we could detect early changes in MRI parameters which would facilitate early diagnosis of prion disease in humans. Such translational biomarkers might become increasingly important to facilitate translation of late-stage potential treatments from animal experiments to humans (see e.g. 11 for a recent review on drug development in PrP). In the RML mouse model we have used, clear evidence of abnormal brain function has been detected about 80 days post-inoculation, when subtle symptoms become apparent in prion-infected mice. Clinically this corresponds to phase 2 of prion propagation where the disease-related prion protein is well characterised throughout the brain (10). MRS measurements which report on neuronal function and metabolism in the brain show deficiencies throughout the disease course. In our experiments, the concentration of Ins, Glu, lactate and lipids were statistically different in all prion-infected mice. Furthermore, MTR abnormalities reporting on macromolecular concentration and composition were found to be reduced at the end-stage of the disease. These findings are discussed in more details below.

### In vivo MRS changes

Changes in brain metabolite concentrations provide key insights about disease progression. Glu is the major excitatory neurotransmitter in the brain. Alterations in Glu concentration are indicative of imbalance in metabolic activity (Krebs Cycle) and a decrease in glutamate concentration is related to a reduced metabolism (12). It is believed that increased concentrations of mI in neurodegenerative diseases reflect an increase of glial cells, because it is primarily expressed in such cells. In prion-infected mice, an increase in mI might be also related to astrogliosis (13).

NAA represents a marker of neuronal density or neuronal function (14). Reduced NAA in prion-infected mice is indicative of neuronal loss, which is similar to evidence given by human studies (15). Other roles of NAA include involvement in lipid synthesis in myelin, both precursor and breakdown product of N-acetylaspartylglutamate, and as storage form of aspartate (16). Under conditions of anaerobic metabolism or inflammation, lactate levels become elevated and a characteristic peak at 1.3ppm overlaps with the lipid resonance (17). In prion-infected mice, our data indicate that both lactate and lipid levels are elevated. To date, limited MRS data have been obtained from individuals with CJD, with similar findings as presented here, in particular showing significant reductions in NAA levels (18), explained by neuronal cell death. Additionally however, a decline in NAA levels might be associated with a functional deficit through synaptic loss.

Changes in Glu, NAA, lactate and lipids were further correlated with astrogliosis in prion-infected mice. There is evidence that astrocytes are the cell type in which the abnormal form of the prion protein is first replicated in the nervous system. It is likely that the changes in astrocytes are due to the “stress” induced on these cells by the misfolded protein. In particular, an *in vitro* study showed that the presence of large numbers of astrocytes can accelerate the rate at which neurons are killed by the toxic peptide (19). Finally, since glial cells increase in number, and because glial cells are thought to produce primarily Lac, which is then used by neurons as primary fluid (20), then this should indeed lead to an increase in extracellular Lac in prion mice, if that increase is not matched by an increase in neuronal metabolism.

### MTR changes

The MTR ratio was found to be significantly increased in 160-day-old mice, compared to that in 80-day-old mice. The increase in MTR could be attributed to decreased water content as a result of the accumulation of myelin, lipids, proteins, proteolipid-proteins, cholesterol and amino-acids. Furthermore, the findings – including MTR changes in healthy mouse brains and reduced MT values in the disease state – are in line with the literature of other neurodegenerative diseases such as Alzheimer’s disease or amyotrophic lateral sclerosis (ALS) (21),(22). Taking this into account, MTR might not likely be a characteristic biomarker of prion diseases as abnormalities have been described in other neurodegenerative disorders. However, in our previous studies, we showed that NOE* is decreased because of a decreased proteasome activity as a result of the accumulation of the aggregated PrP throughout the disease course in the prion-infected group (10) which further correlates with abnormal prion protein deposition and astrogliosis. Therefore, by combining both indices, a useful predictor of clinical disease onset might be established.

### T_1_ and T_2_ relaxation changes

In all regions studied, T_1_ was found significantly different only at the end-stage of the disease in contrary to T_2_ which was significantly higher in thalamus throughout the disease course. These findings support the hypothesis that the early symptoms of the disease are more likely caused by molecular processes which affect normal brain function rather than its macrostructure. A big shortcoming of this study is that we haven’t included any diffusion based microstructural approaches because of the time limitations within of our experiment. Therefore, the extend at which DWI could be useful in prion disease is discussed in reference with existing studies below. It is known that early stages of the pathology are related to synaptic loss resulting into the loss of neurons at the late stages of the disease. Additionally, at the end stage of the disease, extensive vacuolation and brain atrophy alter water distribution which might influence both relaxation times. According to the literature increased *T*_2_ values were observed in prion-infected hamsters suggestive of increased tissue water content which might related to the presence of cell death (23). However, these are qualitative observations taken from T_2_-weighted images, which present with inherent confounds. In contrast, our data were obtained by quantitative methods and provide evidence for widespread degeneration of the brain integrity during the preclinical stages of the disease in this murine model.

### Comparison with other neurodegenerative diseases

Next, we would like to discuss our findings in the context of other neurodegenerative diseases. MT contrast has been used as a novel approach to detect amyloid plaques in the brains of AD mice which are already present in the early stages of the disease. Unlike our results, the MTR was found to be significantly higher in the brains of APP/PS1 and BRI mice, modelling late- and early-stage AD respectively (24). Additionally, MT was found to exceed sensitivity compared to traditional T_2_ measurements. Similar findings were also reported in the Tg2576 mouse model of AD which exhibits increased amyloid beta deposition that eventually progresses into amyloid beta plaque deposition (25). However, in another study using Tg2576 mice quantitative MT failed to pass significance as no alterations in myelin content were observed (26). Interestingly, our results show similarities with patient specific biomarkers observed in AD patients such as hippocampal T_2_ prolongation and MTR reduction as a result of progressive demyelination and reduced capacity of the macromolecules to exchange magnetization with the surrounding water molecules (27).

Other forms of neurodegenerative diseases which are caused by misfolded proteins include Parkinson’s (PD) and Huntington’s disease (HD). In PD mice, the MTR was significantly decreased in the striatum and substantia nigra and it was significantly associated with an increase in the Glu/Cr ratio compared with the control group (28). Additionally, a tendency towards a reduction in T_2_ was measured in the model animals as compared to littermates (29).

MT imaging was used to discriminate symptomatic HD gene carriers from healthy controls and non-affected HD carriers (30), (31). The findings of this study suggest a relationship between disease severity and macromolecular load in caudate nucleus. In our study the absence of plaques or the formation of aggregated proteins at the early stage of the disease does not alter MT which is only relevant at the late stages of prion disease.

^1^H MRS in transgenic HD mice showed a reduction in NAA in the corpus striatum indicating diffuse neuronal loss (32), in other studies this was followed by lower levels of choline and phosphocholine in addition to an increase in glutamine, taurine and myoinositol (33), (34), (35). In accordance with our study, the R6/2 mouse model of HD, also exhibits a decrease in NAA and glutamate (36). In AD mouse models a decrease in NAA and glutamate was observed in addition to an increase in taurine (37). In the APPxPS1 AD model an increase in myo-inositol was also detected (38). Finally, in another study the decrease in NAA and Glu were correlated with the plaque burden (39).

## Conclusion

Several MRI modalities were applied in prion infected mice for gathering information regarding the pathogenesis of this fatal neurodegenerative disorder. Alterations in the brain were detectable by non-invasive quantitative measurements of MRS, MTR, T_1_, and T_2_ values and confirmed by histopathological staining (10). Early signs of the disease were observed at about 130 dpi, however, brain function was already affected from around 80 dpi as shown from our results. It was possible to identify preclinical markers of prion disease for early detection and/or diagnosis. Among all our measurements, decreases in Glu, mI, and elevated lactate and lipids concentration as well as an increase in T_2_ has the potential to warn of impending clinical symptoms. Overall, these results indicate functional down-regulation of brain function in advance of clinical symptom of neurodegeneration. Further studies are required at earlier timepoints for predicting disease onset by means of MR techniques.

## Acknowledgements

We are grateful to MRC technical staff for animal handing during the experiment. We would like to thank Prof Allan Hackshaw and Dr Michael Katsoulis for their input and feedback on mixed effects model analysis. We would like to thank Franscisco Torrealdea and Marilena Rega for insightful discussions and Andreia Silva for assistance during the experiment. This project was funded by UCL Grand Challenge (ED), and by the Department of Health’s NIHR Biomedical Research Centre’s funding scheme to UCL/UCLH (SB, XG, MT). It has also received funding from the European Union’s Horizon 2020 research and innovation programme under grant agreement No 667510 (ED, XG).

